# Engeneering the neurovascular unit: a novel sensorized microfluidic platform to study barrier function and maturation

**DOI:** 10.1101/2025.10.30.685564

**Authors:** Ludovica Montesi, Davide Lattanzi, Matilde Picchi, Stefano Sartini, Tirosh Mekler, Netanel Korin, Rossana Rauti

**Author notes:** These authors contributed equally to this work (Ludovica Montesi, Davide Lattanzi).

## Abstract

Central nervous system diseases pose a significant challenge for the development of effective drugs and therapies. A major limiting factor is the neurovascular unit (NVU), which is both anatomically complex and characterized by a highly selective barrier. Conventional 2D in-vitro models and in-vivo animal models do not adequately replicate its pathophysiology.

Organ-on-a-Chip technology provides a powerful platform to model the NVU, enabling replication of its anatomical and functional features within a dynamic microenvironment that closely mimics the human brain. However, the requirement for specialized facilities and technical expertise limits accessibility, reducing broader translational applications. Additionally, conventional endpoint analyses constrain real-time monitoring of cellular behavior.

Here, we present and validate a novel bi-modular microfluidic chip that offers an easy-to-use and scalable solution for studying cellular cross-talk, while enabling live imaging and real-time measurements. The model incorporates human endothelial cells and primary neurons that were investigated through immunofluorescence and live imaging. The design overcomes key fabrication challenges and integrates a simplified method for Trans-Epithelial/Endothelial Electrical Resistance (TEER) monitoring, allowing in situ real-time assessment of barrier integrity. Overall, this platform represents a robust and versatile tool for in-vitro studies of the NVU, facilitating comprehensive evaluation of its structural and functional dynamics. Our microfluidic NVU-on-chip represents a significant advancement in NVU modelling, providing a versatile platform for CNS drug screening, disease modelling, and personalized medicine applications.

## 1. Introduction

Disorders of the Central Nervous System (CNS) represent the leading cause of disability and the second leading cause of death worldwide (Feigin et al, 2021). One of the major challenges in developing effective treatments for CNS diseases lies in the restrictive barrier imposed by the neurovascular unit (NVU). The NVU is a highly specialized multicellular interface that integrates vascular, glial and neuronal components to maintain brain homeostasis. Its architecture comprises endothelial cells, tightly connected through adherens and tight junctions that strictly regulate permeability (Tietz, Engelhardt, 2015); pericytes, which provide structural and functional support (Uemura et al., 2020); astrocytes, glial cells that mediate neurovascular communication (Abbott et al., 2006); microglia, the resident immune cells of the brain (Thurgur et al., 2019); and neurons, the primary information-processing units of the CNS.

The NVU orchestrates the bidirectional exchange of nutrients, metabolites and drugs between the systemic circulation and the CNS, ensuring a tightly regulated environment for neuronal function (Itoh and Suzuki, 2012). This dynamic interplay between molecular transport and metabolic activity across NVU cell populations is essential for normal brain physiology, and its distruption rapidly leads to neurological dysfunction (Lecrux and Hamel, 2011; Magistretti and Allaman, 2015). However, current experimental models, including animal systems, fail to recapitulate the spatial-temporal complexity of NVU dynamics and the intricate crosstalk among neurons, endothelial cells and the perivascular niche (Vasilopoulou et al., 2016; Nikolakopoulou et al., 2020). Moreover, animal studies are hampered by species-specific differences in efflux transporters, tight junctions and cell–cell signaling (Warren et al., 2009; Nikolakopoulou et al., 2020).

Emerging Organs-on-a-Chips (OoC) technologies offer a promising alternative to overcome these limitations. OoCs are microfluidic *in-vitro* systems engineered to recapitulate the functional units of human organs at a miniaturized scale (Maoz et al., 2021; Monteduro et al., 2023; Guarino et al., 2025). These platforms recreate physiologically relevant tissue microenvironments to enable predictive studies of human biology, disease mechanisms, and therapeutic responses, while reducing reliance on animal testing (Cho et al., 2023; Leung et al., 2022; Nikolakopoulou et al., 2020; Rauti et al., 2021; Rauti et al., 2023).

In the field of neurovascular research, OoCs technology has emerged as a powerful tool to model the human neurovascular unit, whose intricate structural and cellular organization makes it particularly challenging to reproduce compared with other systems (Bhalerao et al., 2020; Caffrey et al., 2021).

In recent years, several *in-vitro* NVU models have been developed to better mimic the physiological conditions, with a focus on replicating the geometry, spatial organization and interactions of cellular components (Lyu et al., 2021; Maoz et al., 2018). Although these platforms represent significant advancements toward faithfully recapitulating *in-vivo* NVU environment, each present limitations that restrict their application. In particular, to date, no single system simulatenously fulfills all the following criteria: modularity, low cost, easy to use, applicable to high-throughput experiments, accurate representation of cell-cell interactions, controlled perfusion capability, and compatibility with high-magnification imaging procedures.

Moreover, integrating non-invasive, real-time sensing technologies to monitor vascular barrier properties remains challenging in most of the NVU-on-chip configurations.

In this study, we present a novel NVU-on-chip model featuring a modular, 3D printed patform designed to meet the demanding criteria outlined above, while enabling real-time monitoring of barrier properties through integrated electrodes. After describing the development of the sensorized microfluidic system, the assembled NVU-on-chip was characterized to validate the platform’s capacity to accurately monitor the formation and maturation of both endothelial and neuronal networks.

Our microfluidic NVU-on-chip represents a significant advancement in NVU modelling, providing a versatile platform for CNS drug screening, disease modelling, and personalized medicine applications.

## 2. Materials and methods

### 2.1 Microfluidic Device Design and Fabrication

The microfluidic platform was designed using CAD Autodesk Fusion 360. The platform is 3D printed in biocompatible resin (BioMedClear V1) with Formlabs Form 3B+ (Formlabs, Somerville, MA, USA). Prior to printing, model surfaces were checked, and supports were added using PreForm software (PreForm 3.0.1, Formlabs, Inc.). After printing, the chips were washed in isopropyl alcohol (Avantor), to remove the unreacted resin, and then cured and dried in a UV curing system (Formlabs).

The OoC is composed of two main blocks connected through a permeable polyethylene membrane (0.4 μm pore size, it4ip S.A., Belgium). The membrane is placed between two cut out Parafilm® (Merck) sheets, thus creating a sealed environment. Each single block is closed by a round coverslip (13 mm Ø), in the lower one for high resolution imaging, while in the upper resin mold for brightfield images. The glass coverslip is sealed to the system with bioresin and UV curing.

### 2.2 Sterilization of the Microfluidic Device

The assembled chip was connected through a system of tubing to a peristaltic pump, and then sterilized with ethanol (5 mL for each chamber), rinsed with MilliQ water (5mL for each chamber), and then washed with Dulbecco’s Modified Eagle Medium (DMEM 1X, Gibco) (5mL for each chamber). Lastly, 30 minutes of UV.

### 2.3 Fluid Computational Model

Computational fluid dynamics (CFD) simulations were conducted using commercial software (Ansys Inc., Fluent® 24R2, 2024) to determine the wall shear stress (WSS) distribution on the PET membrane connecting the upper and bottom parts of the chip, as well as the internal flow field. Due to symmetry, the simulations were performed on only the bottom part of the chip. The computational domain was meshed using 3 million cells with three inflation layers at the walls. The fluid was modeled as incompressible and Newtonian, with properties like water (density ρ = 998.2 kg/m^3^ and dynamic viscosity μ = 0.001 Pa·s). Steady-state simulations were performed with inlet boundary conditions corresponding to flow rates of 50, 100, 150, and 200mL/min, zero-pressure conditions at the outlet, and no-slip conditions at the walls.

### 2.4 Integration of TEER Measurements in the Microfluidic Platform

The barrier properties of the endothelial monolayer were assessed by incorporating a four electrode Transendothelial Electrical Resistance (TEER) system into the OoC. TEER values (Ω/cm^2^) were recorded over time and normalized against the values obtained from an OoC without cells, considered as a blank.

#### 2.4.1 TEER Circuit and Programming

##### Electrical Circuit

The circuit for TEER measurement was implemented using an LM358 operational amplifier interfaced with an Arduino Nano, as shown in **Figure 1**.

**Figure 1.**
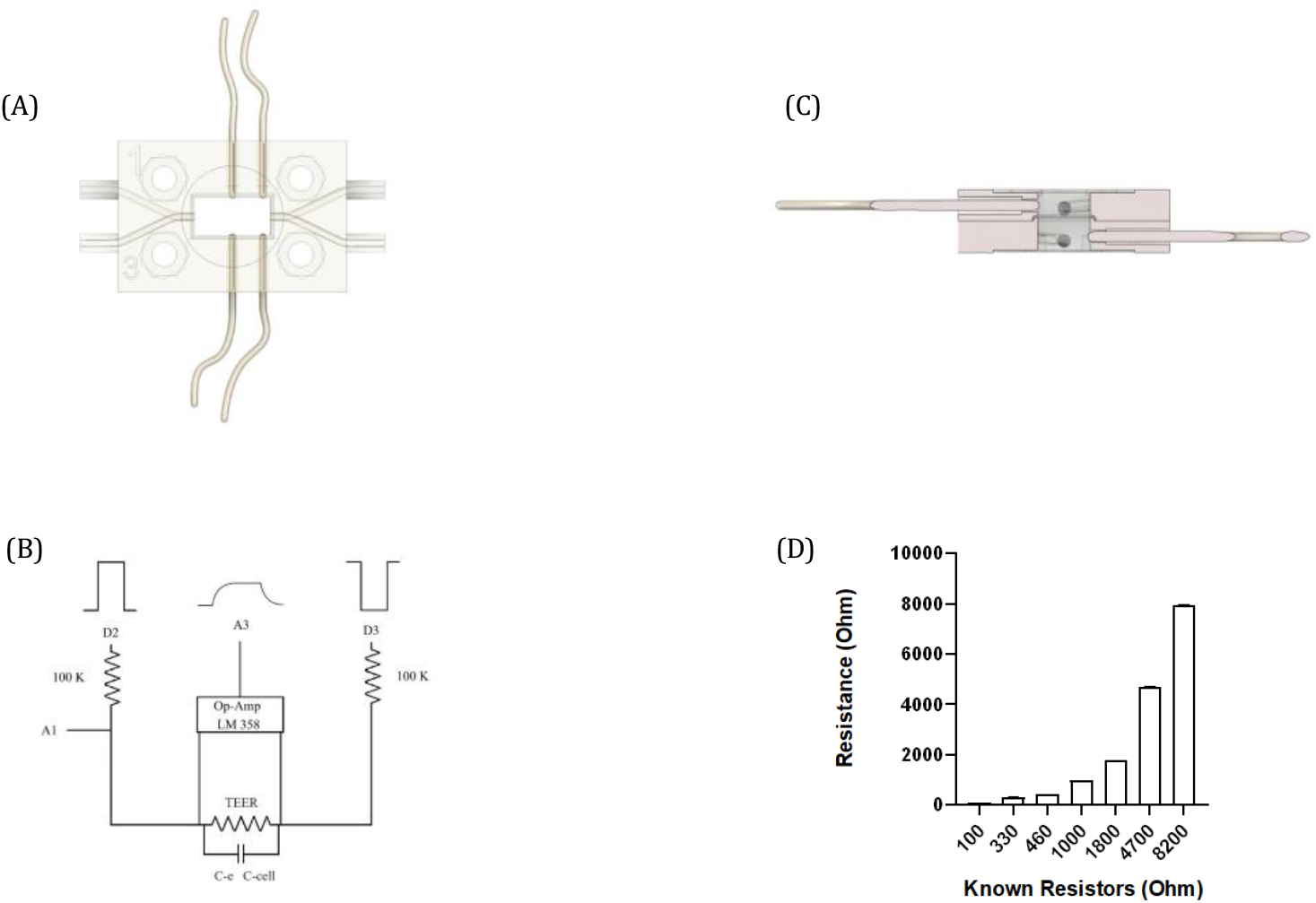
TEER Characterization. (A) Top view of the modular organ-on-a-chip device with integrated TEER measurement electrodes. (B) Cross-sectional view of the chip showing the electrode placement within the dedicated microchannels. (C) Schematic of the custom-built circuit used for TEER acquisition. (D) Calibration plot obtained by measuring a series of standard resistors ranging from 100 to 8200 Ω. Data are expressed as mean ± SE.

The Arduino controls the acquisition and excitation sequence through two analog pins (A1, A3) and two digital outputs (D2, D3) respectively. Pins D2 and D3 are alternately set to high and low logic states to generate a bipolar square-wave excitation across the cell culture, through two 100 kΩ series resistors (Lattanzi et al., 2025). Analog input A1 measures the potential drop across a reference resistor to calculate the current passing through the system. The voltage across the cell monolayer is amplified by the LM358 operational amplifier, and the resulting signal is read by analog input A3.

##### Arduino Board Description

Arduino boards are compact microcontroller platforms widely used in the development of bioelectronic systems. Their analog input pins enable the acquisition of continuous voltage signals from external sensors, which are then converted into digital values by an onboard analog-to-digital converter (ADC). The ADC typically has a 10-bit resolution, producing values from 0 to 1023. With a 5 V reference, the resulting voltage resolution is approximately 0.048 V per step. For improved sensitivity, particularly for low-amplitude signals, the internal reference voltage of 1.1 V can be selected using analogReference(INTERNAL);, enhancing the ADC resolution to about 0.012 V per step. The digital I/O pins can handle a maximum of 40 mA current and 5 V voltage, enabling connection to actuators, LEDs, and other digital devices commonly used in bioelectronic circuits. Programming is conducted through the Arduino Integrated Development Environment (IDE), which uses a simplified version of C++ and offers extensive libraries and example code. This facilitates rapid prototyping and iterative experimentation, making Arduino platforms particularly suitable for bioelectronic research and educational purposes. Among the various Arduino models, the Arduino Uno and Arduino Nano are most frequently employed. The Uno features 14 digital I/O pins, 6 analog inputs, a 16 MHz clock, USB connectivity, and a power jack. The Nano, a smaller form factor, provides 22 digital I/O pins, 8 analog inputs, and is ideal for compact or wearable biosensing devices. In summary, Arduino boards combine flexibility, ease of programming, and precise analog/digital interfacing, making them highly suitable for bioelectronic applications ranging from laboratory prototypes to advanced biomedical research setups.

##### Four Poles TEER Platform Design and Fabrication

A custom-built four-electrode system was developed for the measurement of Trans-Epithelial Electrical Resistance (TEER) in an organ-on-a-chip platform. All electrodes are integrated into the central microfluidic chambers of the organ-on-a-chip to ensure direct electrical contact with the culture medium. The instrument is based on an Arduino Uno microcontroller connected to an LM358 operational amplifier module and a 16×2 I2C LCD display for real-time monitoring. The system measures the voltage drop across a known resistor (100 kΩ) to determine the current (I) and the effective voltage across the sample (Vsample), compensating for the amplifier gain (4.61×) in software. The TEER value (Ω) is then calculated using Ohm’s law:

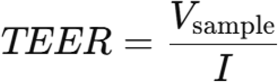

The resulting values for current and TEER are displayed on the LCD screen and transmitted via the serial interface for external data logging and analysis.

##### Detailed Technical Explanation of the Arduino-Based TEER Measurement Code

The bottom presented code performs sequential voltage and current measurements, applies appropriate averaging and scaling factors, and outputs the computed TEER values both to a 16×2 I2C LCD display and to the serial monitor for data logging.

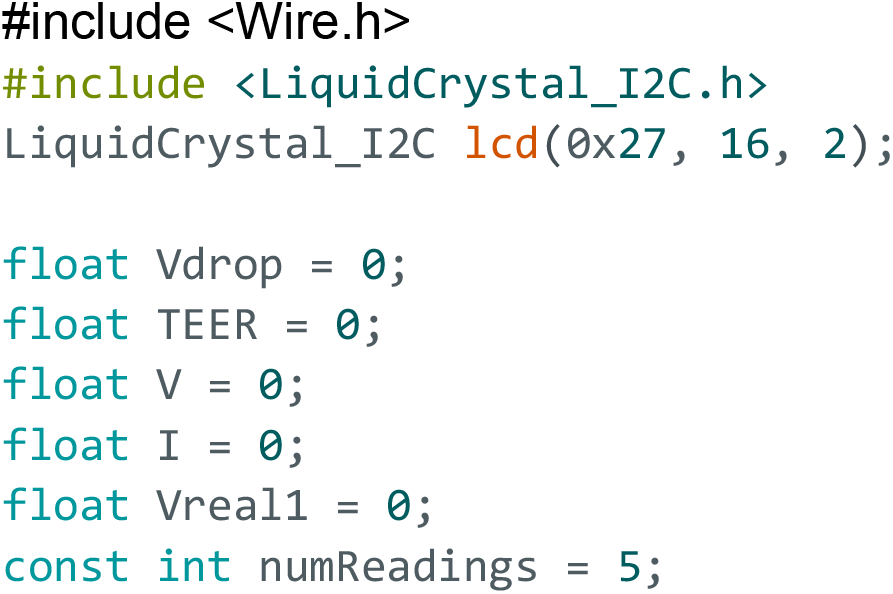
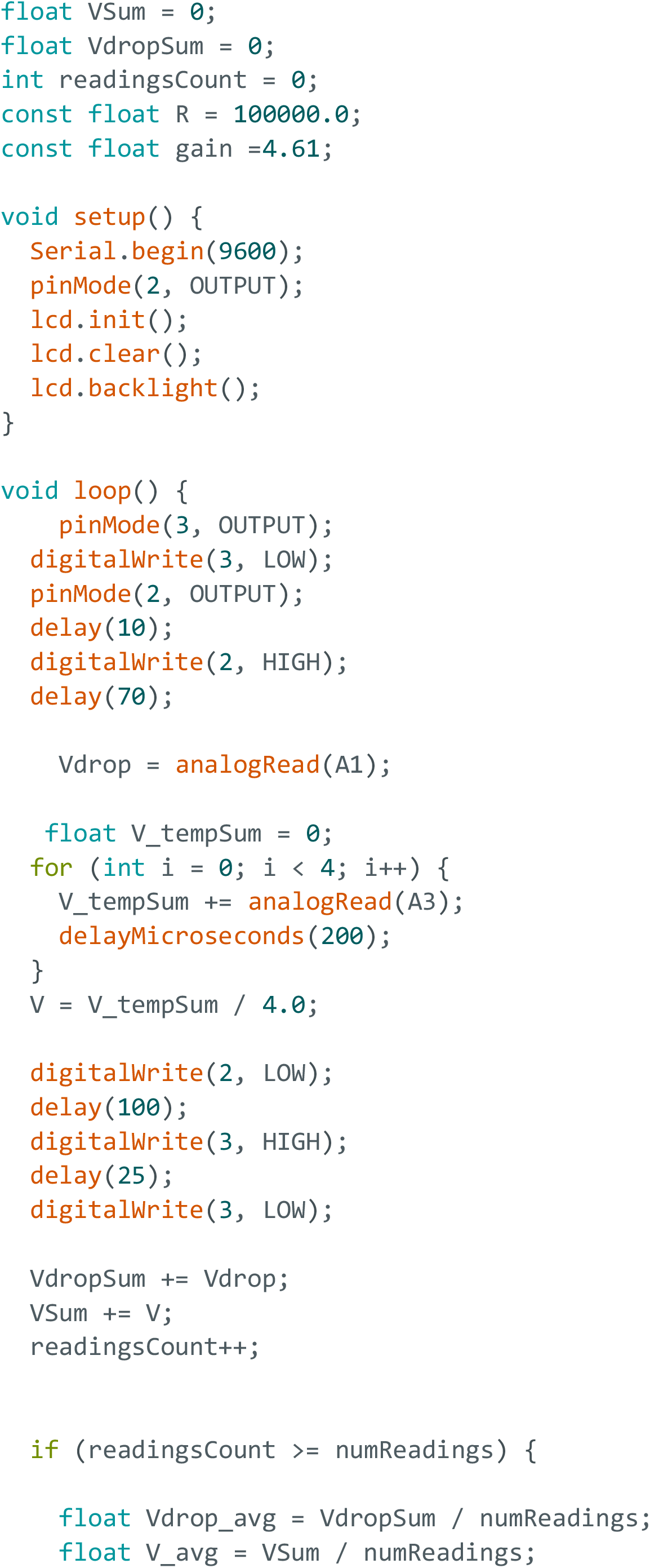
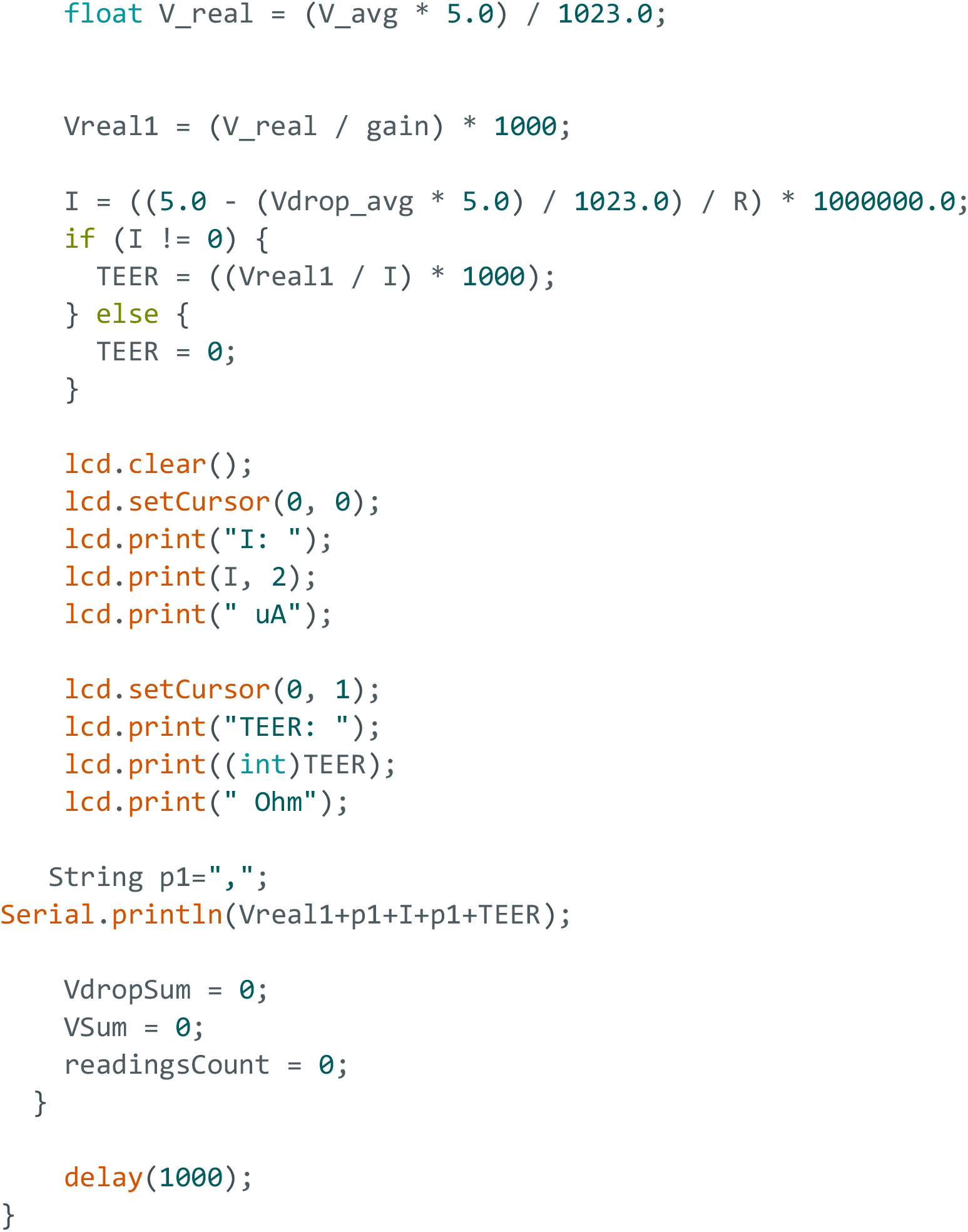

##### Hardware Configuration and Initialization

At the beginning of the program, the LiquidCrystal_I2C library is initialized with the I2C address 0x27, corresponding to a 16×2 LCD module. This module is used to display real-time measurement data. The setup routine initializes serial communication at 9600 baud, configures the necessary digital pins, and activates the LCD backlight. A known resistor of 100 kΩ in series with D2 digital pin is employed as a reference element for current calculations, while the LM358 amplifier provides a voltage gain of approximately 4.61, compensating for small signal levels in the measurement circuit.

##### Measurement and Sampling Routine

In the measurement routine, the D2 and D3 pins are alternately configured as either output or ground, being set to a HIGH or LOW logical state in succession. This alternating configuration enables the generation of a bipolar excitation signal across the sample, ensuring symmetrical current flow through the cellular monolayer. The approach reproduces the stimulation strategy described by Lattanzi et al. (2025), improving measurement stability and minimizing electrode polarization effects. Furthermore, the 100 kΩ resistors connected in series with D2 and D3 serve a dual purpose: they function as reference elements for current determination and act as current-limiting resistors, restricting the current delivered to the cells to physiologically safe levels. This configuration prevents potential damage to the biological sample during TEER measurements while maintaining the precision and reproducibility of the electrical recordings. Analog voltage readings are then obtained from the analog input pins: A1 measures the voltage drop across the known 100 kΩ resistor, which is used to calculate the current flowing through the circuit; A3 measures the potential difference detected by the voltage electrodes across the sample, amplified by LM358. This process is repeated five times, and the corresponding voltage readings are accumulated to compute mean values (Vdrop_avg and V_avg) over multiple iterations. Averaging over several readings increases the measurement stability and reduces the influence of stochastic noise or transient signal variations.

##### Data Conversion and Calculation

Raw analog values, ranging from 0 to 1023, are converted to volts using the standard Arduino ADC conversion factor for a 5 V reference (V = (raw * 5.0) / 1023). The resulting total voltage (V_real) is further divided by the amplifier gain to obtain the true potential difference across the biological or electrical sample (Vreal1), expressed in millivolts.

The current (I) through the system is computed from Ohm’s law using the known reference resistor (R = 100 kΩ). Specifically, the code determines the voltage drop across this resistor and converts it into microamperes:

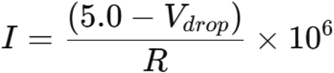

Once the current and the effective voltage across the sample are known, the TEER value is obtained as:

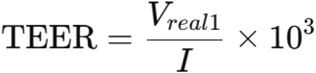

This calculation provides the electrical resistance of the epithelial and membrane barrier in ohms. A conditional statement ensures that division by zero is avoided in the event of negligible current flow.

##### Data Output and Visualization

The computed parameters are displayed on the LCD in real time, showing the current (in microamperes) and the TEER value (in ohms). Simultaneously, the same data are transmitted via the serial interface for external recording or further analysis. After each complete averaging cycle, the temporary variables are reset, and the loop continues with a one-second delay between successive measurement cycles.

In summary, this code implements an automated, low-noise TEER measurement system based on the Arduino Uno platform. It integrates analog signal amplification, digital sampling, data averaging, and real-time display functionalities. The use of averaging routines and amplifier compensation enhances accuracy and stability.

### 2.5 Cell Cultures

#### Endothelial culture

For the endothelial model, Human Umbilical Vein Endothelial Cells (HUVEC, PromoCell) were used. After thawing, the HUVECs were expanded in a low-serum endothelial cell growth medium (Promocell), at 37 °C with 5% CO_2_ in a humidifying incubator, and used at passage p2-p5. Cells were grown at 80-90% confluence before being seeded inside the microfluidic platform. Before seeding, the PET membrane was treated with Entactin-Collagen IV-Laminin (ECL) Cell Attachment Matrix (Merck) diluted in Dulbecco’s Modified Eagle Medium (DMEM; 10 μg/cm^2^), for 1 h in the incubator. Then, the HUVECs, harvested using trypsin/EDTA solution, were seeded inside the chip at a density of 20.000/cm^2^. After 1 h at 37 °C with 5% CO_2_ in a humidifying incubator, the flow was activated at 6 µL/min.

#### Neuronal culture

Primary dissociated neurons were obtained from the spinal cord of 7 days old chick embryos. According to the Directive 2010/63/EU, chick embryos up to embryonic day 14 are not considered live vertebrate animals and are therefore not subject to animal experimentation regulations. Prior to cell seeding, the glass coverslip was treated with poly-L-ornithine (0.01%, Sigma-Aldrich) overnight at 4 °C. Then, the substrates were rinsed three times with sterile MilliQ water, and treated with laminin (1.4 mg/mL, Sigma-Aldrich) diluted in sterile MilliQ water (10 μg/mL) for 1h at 37° with 5% CO2 in a humidifying incubator. Cells were seeded at 300.000 cells/cm^2^ in Neurobasal Medium (Gibco), supplemented with FBS (10%, Gibco), B27 (2%, Gibco), Glutamax (1%, Gibco), and PSA (1%, Gibco), and Gentamicin (1%, Sigma-Aldrich). After 1 h at 37 °C with 5% CO_2_ in a humidifying incubator, the flow was activated at 6 µL/min. Media change occurred every other day, until day 7.

### 2.6 Immunofluorescence Analysis

HUVEC cells and primary neurons cells were rinsed in Phosphate Buffered Saline (PBS) and fixed in 4% paraformaldehyde for 20 min at Room Temperature (RT). Immunocytochemistry was carried out after permeabilization with 0.1% Triton X-100 (Sigma-Aldrich) in PBS for 10 minutes RT and blocking for 30 min with 5% FBS in PBS. The primary antibodies, anti β-catenin (1:200; Cell Signaling), β-tubulin III (1:200, Merck), and anti-Glial Fibrillary Acidic Protein (GFAP, 1:400; Cell Signaling) were applied overnight in PBS and 5% FBS, at 4 °C. Cells were then washed in PBS three times and then incubated with the secondary antibody, anti-rabbit Alexa Fluor-594 (1:500; Invitrogen) or or anti-mouse Alexa Fluor-488 (1:500; Invitrogen) in 5% FBS in PBS for 2 hours at RT. After two washes with PBS, cells were incubated with DAPI (1:10.000 in PBS; Cell Signaling) for 10 min at RT to stain the nuclei. After two washes in PBS, imaging was carried out using an inverted epifluorescence microscope (Nikon Eclipse Ti2), with appropriate filter cubes and equipped with 20x/0.50 NA and 40x/0.75 NA objectives. Image reconstruction and processing were done using the open-source imaging software Fiji (Imagej; Schindelin et al., 2012).

### 2.7 Live Calcium Imaging

For Calcium (Ca^2+^) imaging experiments, primary neuronal cultures were loaded with the cell-permeable Ca^2+^ dye Oregon Green 488 BAPTA-1 AM (Life Technologies). Cultures were incubated for 45 minutes at 37° with 5% CO_2._ Subsequently the media was rinsed, and replaced with a recording solution containing (in mM): 150 NaCl, 4 KCl, 1 CaCl_2_, 10 Hepes, 10 glucose (pH 7.4). The NVU-chip containing the Oregon Green loaded cultures was placed on an inverted microscope (Nikon Eclipse Ti2) and observed with a 20x objective (0.50 NA) using a DS-10 Camera (Nikon). Images were acquired every 250 ms, for 5 minutes, under continuous illumination. The Ca^2+^ dye was excited at 488 nm using appropriate filters cube set and a LED lamp (Nikon). Video analysis was done using the open-source imaging software Fiji (Imagej; Schindelin et al., 2012), and Clampfit software (pClamp suite, 10.2 version; Axon Instruments).

### 2.8 Barrier Disruption

EGTA (ethylene glycol-bis(β-aminoethyl ether)-N,N,N′,N′-tetraacetic acid; Sigma-Aldrich) was prepared at 5 mM with dH2O; pH was titrated to 7.4 with HCl 1 M. The medium in the bubble trap was entirely replaced with the solution containing EGTA and TEER was measured after 50, 70, 75,120,135 and 150 min.

### 2.9 Statistical Analisys

TEER (Ω cm^2^) values were recorded from ten consecutive measurements in the chip each day. For each measurement, the mean value recorded on day 0 (considered as the blank) was subtracted to normalize the data. Daily averages were then calculated, and results are presented as the average of every two days. All graphs were generated using GraphPad 8 (GraphPad Prism Software).

## 3. RESULTS

### 3.1 Microfluidic NVU-on-Chip Model

The NVU-on-Chip model developed in this study consists of a microfluidic chip with two compartments separated by a porous, transparent Polyethylene (PET) membrane (**Figure 2**), designed to mimic the *in-vivo* microenvironment of the NVU. As described in the Materials and Methods section, the chip is made of biocompatible transparent resin. The device comprises an upper chamber respresenting the vascular compartment and a lower chamber representing the parenchymal (neurons and astrocytes) compartment. Each module measures 30 mm in length, 20 mm in height, and 3 mm in depth, and features a central rectangular chamber measuring 10 mm x 5 mm (length x height). Two esternal channels (5 mm long, 1.3 mm internal diameter, 2.5 mm external diameter) allow connection to a peristaltic pump (**Figure 2**). The external faces of the modules include a circular groove (13.4 mm in diameter) to improve the adhesion of the coverglass, and four corner holes (3.6 mm in diameter) for bolts (3 mm diameter, 6 mm length) and nuts to secure the assembly. Once the entire device is sealed, circular coverslips are glued to the outer faces using the same resin as the modules (**Figure 2A, 2B and 2C**). Tygon tubing connects the chip channels to the peristaltic pump, and additional tubing connects the system to a bubble trap that removes air bubbles and serves as a liquid reservoir. This configuration allows independent flow control in each chamber of the chip **(Figure 2D)**.

**Figure 2.**
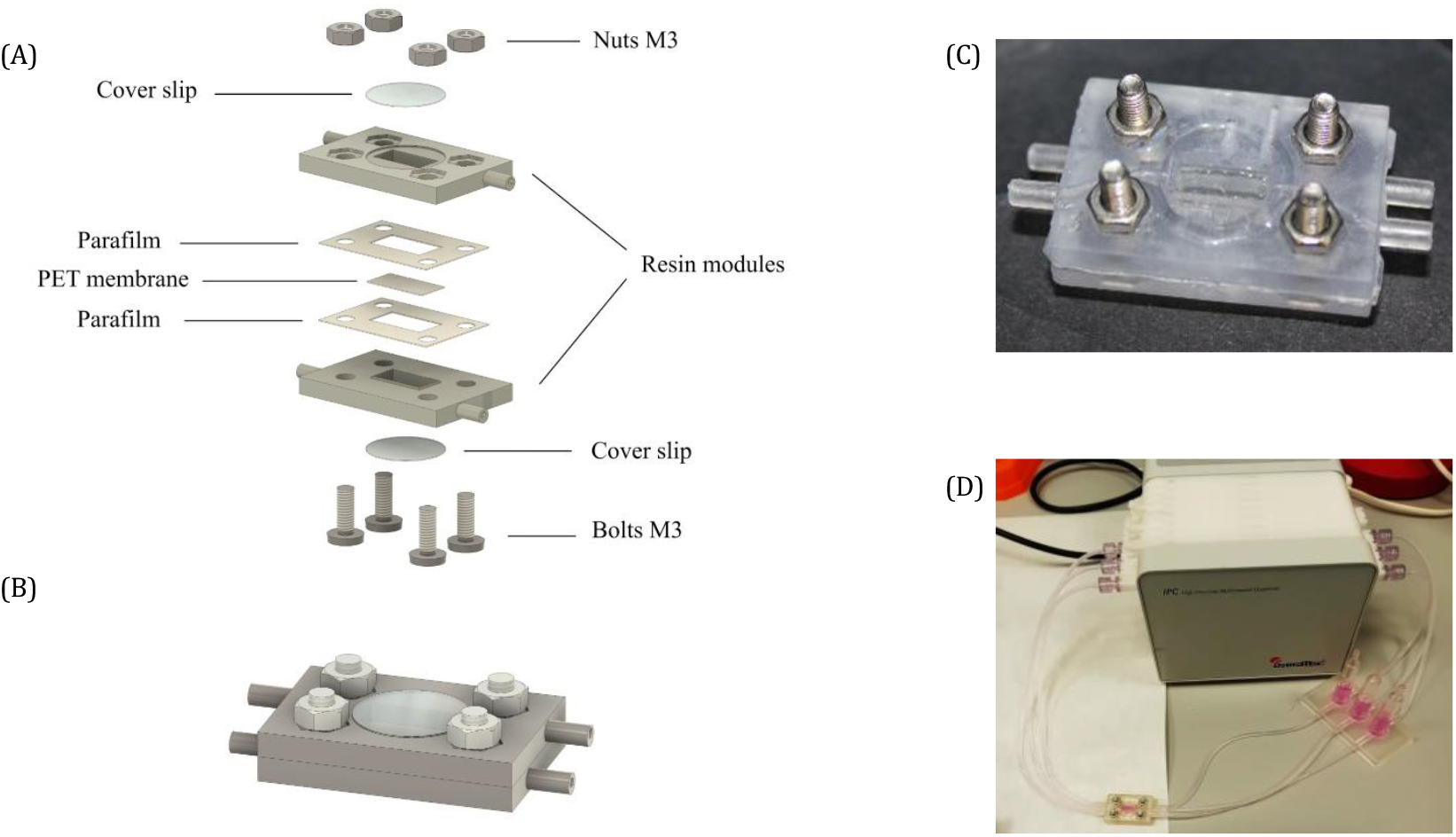
NVU-on-a-Chip Design. (A) Exploded view of the chip, showing the assembly from bottom to top: i) bolts, ii) coverslip, iii) resin mold, iv) parafilm, v) PET membrane, vi) parafilm, vii) resin module, viii) coverslip, and ix) nuts. (B) 3D illustration of the fully assembled NVU-chip. (C) Photograph of the assembled chip made of biocompatible transparent resin. (D) Photograph of the NVU-chip connected to a peristaltic pump via Tygon tubing and bubble trap.

### 3.2 Computational simulation

**Figure 3A** illustrates the velocity streamlines inside the model for all tested flow rates. It can be observed that, within the rectangular domain immediately beneath the PET membrane, the velocity decreases significantly and remains relatively constant across all flow conditions. **Figure 3B** presents the Wall Shear Stress (WSS) distribution, indicating higher WSS values at the central region of the model, while relatively low WSS values occur near the inlet and outlet of the rectangular domain across all flow rates tested.

**Figure 3.**
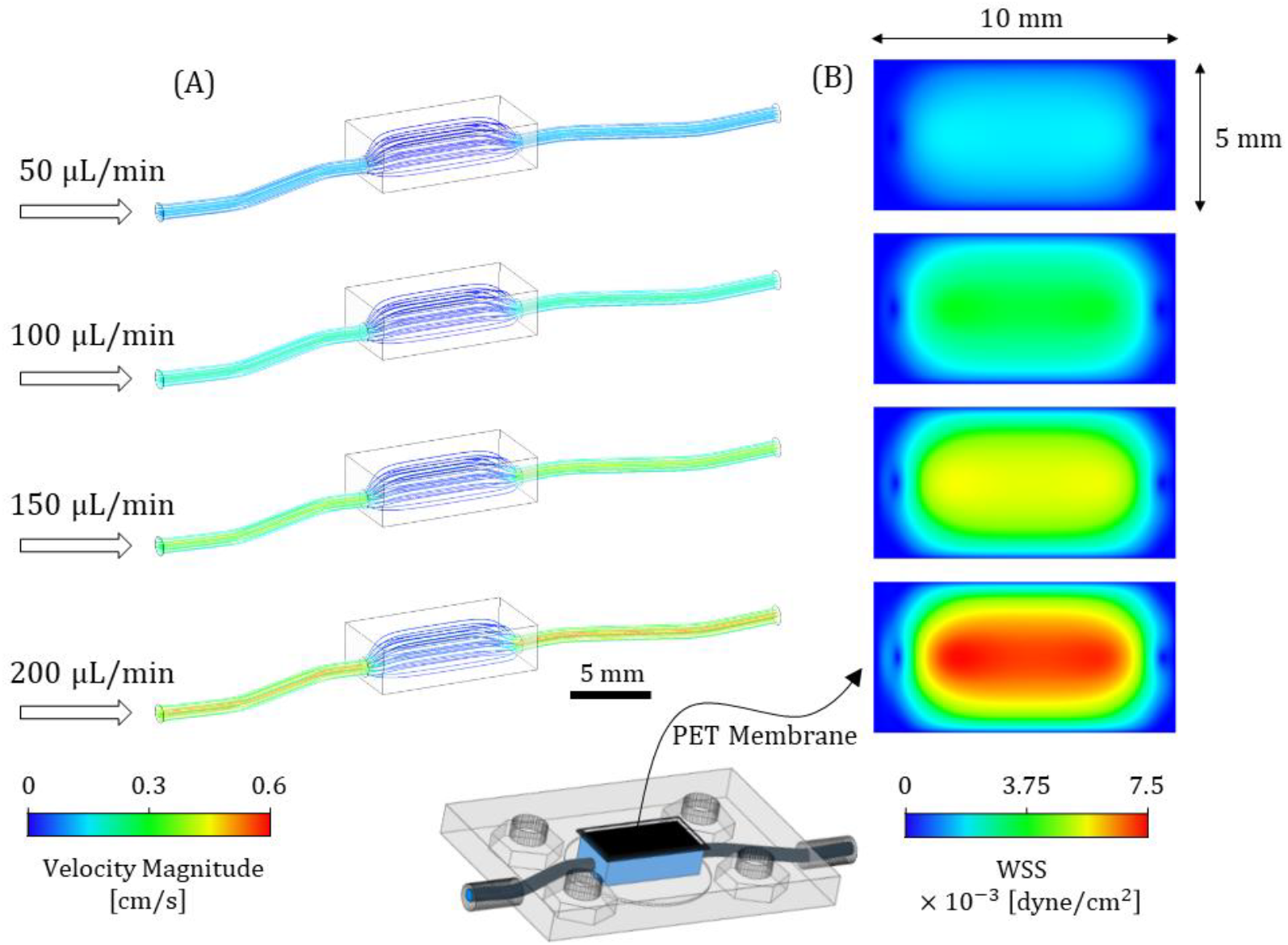
**Computational fluid dynamics (CFD) simulations** of the bottom part of the NVU-chip for the four tested flow rates: 50, 100, 150, and 200 μL/min. (A) Pathlines illustrating velocity magnitude for all flow rates tested. (B) Wall shear stress (WSS) distribution on the PET membrane connecting the bottom and upper parts of the NVU-chip.

### 3.3 Formation of the NVU Model

To reproduce the cerebral neurovascular unit (NVU), endothelial cells were cultured in the upper compartment of the device, while neurons and astrocytes were cultured in the lower compartment (**Figure 4**). Cellular adhesion and survival were evaluated to assess device biocompatibility.

**Figure 4.**
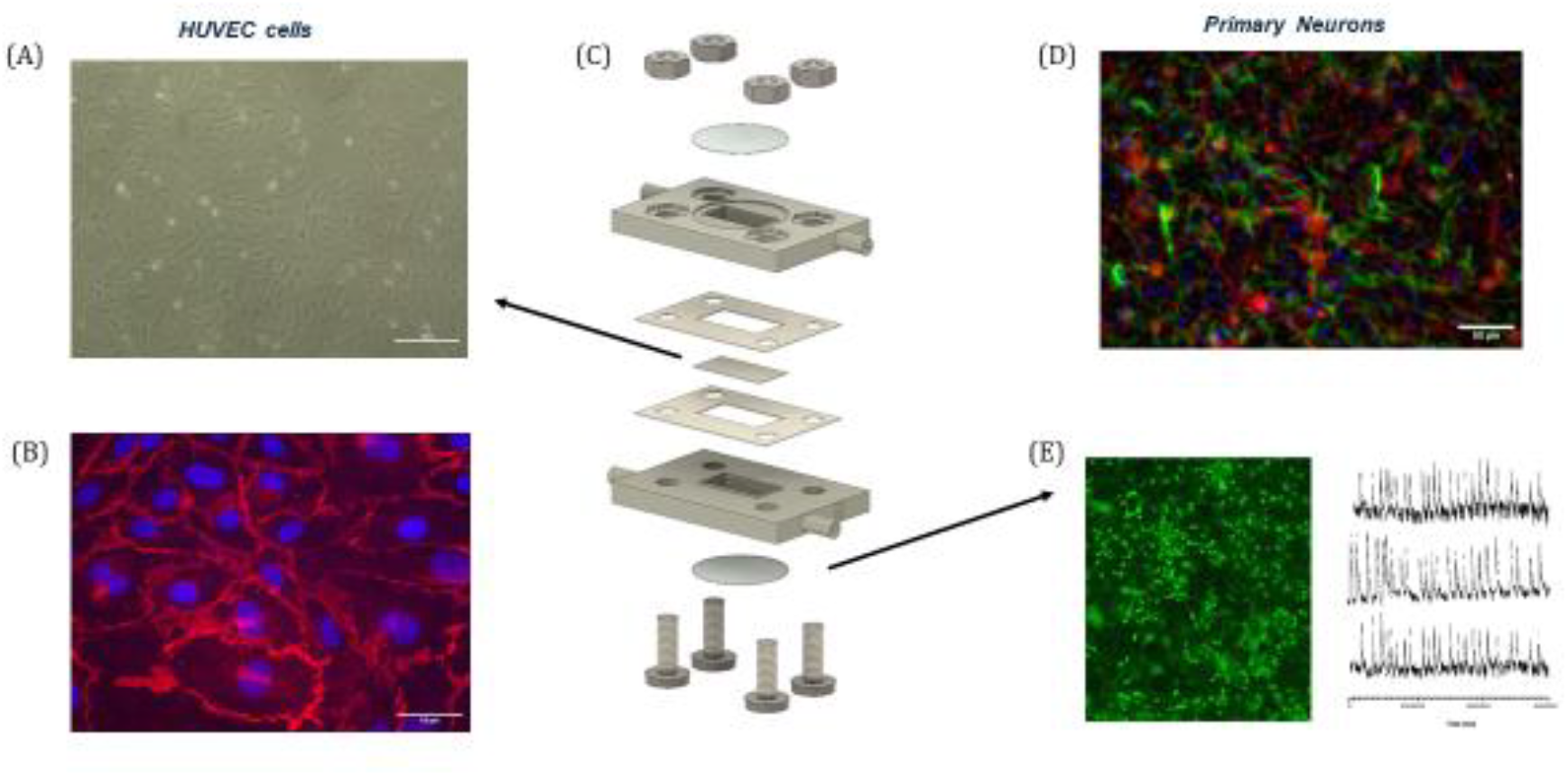
Endothelial and neuronal cell cultures in the NVU-chip. (A) Bright-fied image of HUVEC cells cultured within the NVU-chip. (B) Epifluorescence image reconstruction of HUVECs stained for VE-Cadherin (red) and DAPI (blue). (C) Exploded schematic view of the NVU-chip structure. (D) Epifluoresecnce reconstruction of primary neurons cultured in the NVU-chip, stained for β-tubulin (red), GFAP (green) and DAPI (blue). (E) Epifluorescence image of primary neurons labeled with the calcium indicator Oregon green 488, and representative calcium transients recorded from primary neurons in the NVU-chip.

The two cell types were seeded on different days: neurons were plated immediately after dissociation from chick spinal cords, whereas HUVEC endothelial cells were seeded the following day. One of the advantages of this system is the flexibility to culture cell types separately, allowing each to mature separately and under optimal conditions. This approach ensures that all cell populations reach functional maturity simultaneously, enabling the development of a physiologically relevant NVU model. After 6 days in culture, cell arrangement, adhesion, and maturation were assessed by immunofluorescence staining for VE-Cadherin, a key tight junction protein expressed by endothelial cells, β-tubulin, a neuronal marker, and GFAP, a glial marker (**Figure 4 B and 4D**). The endothelial layer displayed uniform adhesion across the entire surface (**Figure 4A and 4B**) and clear VE-Cadherin expression (in red, **Figure 3B**), indicating the formation of functional tight junctions. Similarly, by stayining neurons for β-tubulin (in red) and glial cells for GFAP (green) we demostrated their survival and maintenance of neuroglia identity within the NVU-on-Chip **(Figure 4D**). To further investigate neuronal network dynamics, calcium imaging was performed using the fluorescent indicator Oregon Green 488 BAPTA-1 AM. This minimally invasive technique enables real-time monitoring of calcium transients at single-cell resolution, providing insights into neuronal activity within the microfluidic NVU-chip (**Figure 4E**).

### 3.4 In Situ Monitoring of Barrier Formation and Maturation

A key limitation of current NVU-chip models is the lack of in situ, real-time analytical tools, as most approaches rely on end-point measurements that prevent dynamic assessment of barrier functionality. Several techniques and molecular tracers have been developed to evaluate permeability; however, Transendothelial Electrical Resistance (TEER) remains the most rapid and non-invasive method for real-time measurements. TEER quantifies the electrical resistance across a cellular monolayer by applying a small current between electrodes and measuring the resulting voltage (Malik et al., 2025). Although widely used in conventional cell culture systems to assess barrier integrity, integrating electrodes into OoC devices is particularly challenging. The compact and sub-millimetric size of microfluidic channels restrict direct access to the cellular layer, complicating TEER acquisition (Henry et al., 2017). The modularity of our OoC platform enables customizable electrode integration. Specifically, we modified the CAD design to include lateral channels extending from the device exterior to the inner chamber, allowing the insertion of electrodes **(Figure 1 A-B)**. This configuration permits TEER measurements across the cellular monolayer cultured on the central membrane. We implemented a four-electrode platinum system sealed with the same resin used for module fabrication. Electrodes were connected to an Arduino-based circuit, and TEER values were calculated using Ohm’s law (Lattanzi et al, 2025).

#### 3.4.1 LM358 Operational Amplifier Module

Accurate TEER measurements require minimal excitation currents to preserve cell integrity, resulting in low-voltage signals (in the millivolt range) at the sensing electrodes. The Arduino’s internal voltage reference did not provide sufficient accuracy or stability for these weak signals. An LM358 operational amplifier was therefore integrated into the circuit to amplify the voltage drop across the sensing electrodes. The LM358 increases the signal amplitude to a range detectable by the Arduino’s ADC while maintaining signal fidelity. Its low noise, high input impedance, and low-voltage operation make it suitable for microcontroller-based acquisition.

This amplification stage enables reliable real-time TEER measurements by (i) maintaining safe current levels for cell viability and (ii) improving the signal-to-noise ratio for precise resistance calculation.

#### 3.4.2 Importance of the Four-Electrode Configuration

The four-electrode (four-terminal) configuration is critical for accurate and reproducible TEER measurements, especially in biological and microfluidic systems where electrode polarization and solution resistance can introduce significant artefacts. In conventional two-electrode setups, the same electrode pair is used for both current injection and voltage sensing, leading to measurement errors due to electrode– electrolyte interface impedance, polarization effects, and surface drift, especially in ionic culture media and during long-term experiments. In contrast, the four-electrode configuration separates current injection from voltage measurement. Two electrodes apply the excitation current through the sample, while the remaining pair senses the potential drop across the cellular barrier without drawing any significant current. This separation minimizes artefacts arising from electrode impedance and ensure stable, high-fidelity TEER measurements over time.

Because the voltage-sensing electrodes operate under near-zero current conditions, their polarization is minimized, and the measured voltage accurately reflects the true trans-epithelial potential difference, independent of contact impedance and medium resistance. This configuration is particularly advantageous in organ-on-a-chip systems, where microchannel geometries and limited electrode spacing can exacerbate errors arising from electrode degradation or medium conductivity changes. Platinum electrodes were selected for their chemical inertness, low corrosion rate, and high stability in physiologically relevant media containing salts, proteins, and metabolites. Over time, platinum minimizes electrochemical reactions that could otherwise lead to electrode fouling or ion accumulation, both of which distort TEER readings in two-electrode systems. In our system, we further reduce electrode polarization by applying bipolar stimulation to the current-injecting electrodes. This approach, previously described in our work (Lattanzi et al., 2025), reverses the current periodically during each measurement cycle, preventing long-term charge accumulation and Faradaic reactions at the electrode–electrolyte interface. This method was tested in a two-electrode prototype for TEER measurement and has proven effective in maintaining electrode stability over extended experiments.

The four-electrode configuration therefore ensures:

- Improved accuracy, by eliminating contributions from electrode polarization and medium resistance.
- Enhanced temporal stability, allowing long-term monitoring of barrier integrity under dynamic conditions.
- Reduced signal drift, since sensing electrodes are not affected by Faradaic processes.
- Greater reproducibility, particularly in microfluidic devices with limited electrode spacing and small volumes.

Overall, the four-electrode TEER configuration provides a robust and reliable approach for quantifying epithelial and endothelial barrier function in microphysiological systems, ensuring that the measured resistance corresponds predominantly to the cell layer itself, rather than to artifacts arising from the measurement system.

#### 3.4.3 Electrode Configuration within the NVU-on-Chip

The NVU-on-chip device consists of two superimposed microfluidic chambers, an upper and a lower compartment separated by a porous PET membrane (pore diameter 0.4 μm). The membrane supports the formation of a confluent cellular monolayer and enables controlled exchange of solutes between the two chambers, reproducing the architecture and barrier function of endothelial interfaces. For TEER measurements, four platinum electrodes were integrated into the chip following a four-pole configuration. Each compartment contains one current-injecting and one voltage-sensing electrode. In the upper chamber, both electrodes are positioned on the right lateral wall, while in the lower chamber the corresponding pair is placed on the left-hand wall **(Figure 1A-B)**. This staggered alignment ensures a homogeneous current distribution across the membrane and maximizes the effective area of the cellular barrier exposed to the electric field. The electrodes, with a rectangular cross-section of 1.0 mm × 0.5 mm, are mounted flush with the inner wall of each microfluidic chamber to avoid interfecence with fluid flow or direct contact with the cultured cells. This configuration minimizes physical perturbation of the biological sample and prevents localized cell damage or detachment. The selected electrode size provides a balance between low impedance and minimal polarization, maintaining signal stability and reproducibility during long-term experiments. Lateral electrodes placement also reduces capacitive coupling between current and voltage pairs, ensuring a stable and reproducible electrical interface with the culture medium without altering flow dynamics. Overall, this configuration enables non-invasive, and spatially uniform TEER measurements.

#### 3.4.3 TEER Device Validation and Characterization

Before performing biological measurements, the operation of the custom-built TEER measurement system was simulated using the Tinkercad Circuits online platform. The Arduino sketch and the electronic circuit were virtually tested by connecting resistive loads ranging from 0.1 to 10 kΩ in parallel with capacitors between 1 and 1000 nF. These simulations were used to optimize the duration of the bipolar square-wave stimulus generated by the Arduino’s digital outputs, ensuring that the capacitive component of the circuit reached full charge before current measurements were taken. The signal recorded at analog channel A3 exhibited the expected exponential charging and discharging behavior typical of a parallel RC network, reaching a steady-state plateau once the capacitor was fully charged (**Figure 4C**). Extending the square-wave duration was necessary for higher capacitance values to achieve a stable plateau response, with 70 ms proving sufficient for capacitors up to 1500 nF. Following the simulations, the circuit was assembled and tested on a breadboard using resistors and capacitors identical to those employed in the virtual model. The experimental data confirmed the accuracy of the measurement system, with measured resistance values closely matching nominal ones (100 Ω: 0.093 ± 0.2 Ω; 330 Ω: 326 ± 0.9 Ω; 460 Ω: 455.8 ± 0.13 Ω; 1000 Ω: 1003.1 ± 0.64 Ω; 1800 Ω: 1813.5 ± 0.56 Ω; 4700 Ω: 4701.9 ± 2.07 Ω; 8200 Ω: 7973.9 ± 1.01 Ω Fig. 4D). The validated circuit was then integrated into the NVU-chip. To verify the stability of the electrical readings, long-term measurements of the culture medium resistance were performed over several days. The application of bipolar square-wave stimuli stabilized the resistance values within a few minutes, and the reading remained constant throughout the recording period. This confirmed that bipolar stimulation effectively prevents electrode polarization and allows continuous and stable TEER monitoring.

#### 3.4.5 TEER Measurements to Characterize Barrier Functionality

After establishing the TEER setup, we sought to demostrated its potential and validation for assessing endothelilal barrier functionality in the NVU-Chip **(Figure 5 A-B)**. To test the system, we cultured HUVEC cells in the chip under continuos flow conditions, and TEER was monitored daily starting from the day of seeding (day 0), considered as blank. During each measurement, TEER values were obtained by averaging 10 consecutive values displayed on the LCD display. After 14 DIV, the electrical resistace progressively increased up to 950 Ω /cm^2^ (**Figure 5C**). To assess the system’s responsiveness to barrier distruption, we applied EGTA at 5.5 mM in the endothelial compartment. TEER was recorded at 50, 70, 75,120,135 and 150 min after EGTA application (**Figure 5D**). The resistance decrased after 50 min reaching the lower value, comparable to the blank, after 135 minutes, confirming the sensitivity and reliability of the integrated electrodes in detecting dynamic changes in barrier integrity.

**Figure 5.**
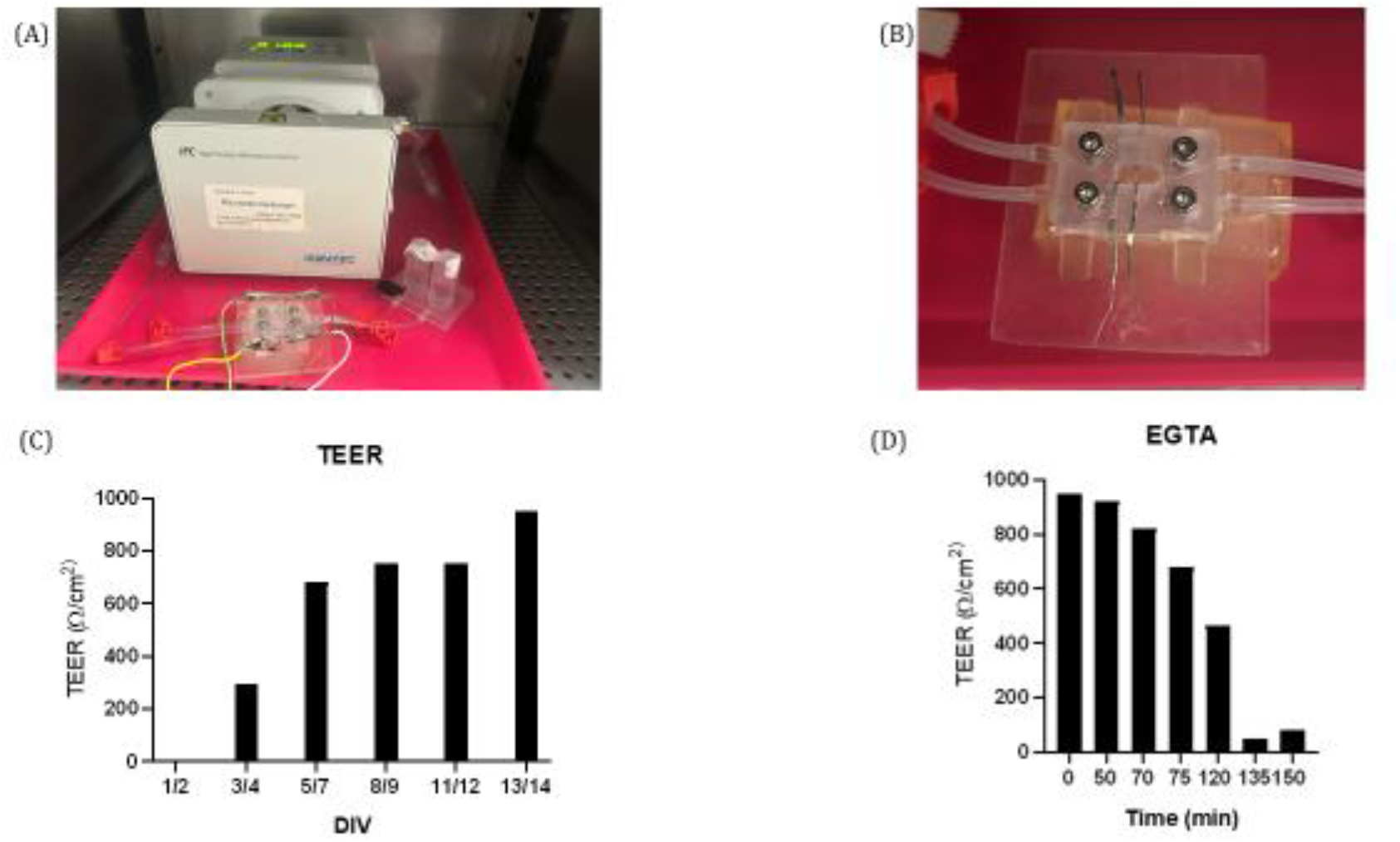
TEER measurements in the NVU-chip. (A) Photograph of the NVU-chip connected to peristaltic pump and TEER system. (B) Photograph of the electrodes inserted in the NVU-chip. (C) Plots showing TEER measurements of HUVEC cells in NVU-chip at different time-points. (D) Changes in endothelial barrier function as a result of 5 mM EGTA drug application.

## 4. DISCUSSION

We present a sensorized NVU-on-chip platform for real-time monitoring of neurovascular cell maturation and barrier integrity, addressing key challenges in CNS drug discovery. By co-culturing endothelial, neuronal and glial cells under dynamic flow, the device recapitulates the *in-vivo* NVU microenvironment, providing a modular, physiologically relevant model superior to conventional static systems.

The microfluidic chip, made from biocompatible resin, integrates platinum electrodes in a four-pole configuration for continuous, non-invasive TEER measurement. TEER progressively increased over time, confirming endothelial barrier formation, while EGTA-induced disruption validated sensitivity. Simultaneous monitoring of neuronal calcium dynamics highlights potential for studying neurovascular coupling and functional CNS responses.

Its modular architecture allows easy expansion with additional membranes or stacked modules for complex tissue interfaces, and multiple chips can be linked in series. Integration of patient-derived cells supports personalized medicine applications.

Conventional TEER methods using wire or external electrodes can compromise sterility or disturb flow, and distant electrodes reduce sensitivity. Our low-cost, custom four-electrode system is manually positioned and fixed with UV-curable resin, ensuring stable, reproducible, and sterile measurements without affecting microfluidic flow. Compared to microfabricated Au or metal-oxide electrodes, which require clean-room fabrication and complex lithography, our approach provides high-performance TEER monitoring with minimal infrastructure.

Overall, this accessible, scalable NVU-on-chip platform enables dynamic assessment of endothelial barrier function and neurovascular activity. Its versatility and modularity make it a robust tool for CNS drug screening, disease modeling, and personalized medicine, bridging the gap between in vitro models and physiological relevance.

## 5. CONCLUSIONS

The proposed sensorized microfluidic device represents a significant advancement in NVU modeling, offering real-time, non-invasive monitoring of endothelial and neuronal integrity and permeability in physiologically and pathologically relevant environments. This platform holds great promise for drug screening, disease modeling, and personalized medicine, providing a valuable tool for advancing our understanding of the NVU and developing new therapies for CNS disorders.

## Credit authorship contribution statement

LM designed and conceived the platform and performed cell biology and immunofluorescence experiments and analysis. DL contributed to chip design and development as well as electrodes integration in the platform. MP contributed to chip design. SS performed primary neurons preparations. TM and NK performed computational simulations. RR conceived the study and the experimental design. LM, DL, SS, MP and RR wrote the manuscript.

## Declaration of competing interest

The authors declare that they have no known competing financial interests or personal relationships that could have appeared to influence the work reported in this paper.

## Acknowledgements

This work was supported by the European Union - NextGenerationEU within the framework of PNRR Mission 4 - Component 2 - Investment 1.1 under the Italian Ministry of University and Research (MUR) programme &quot;PRIN 2022 PNRR&quot; - grant number P2022TKL5T_001 Calinero - CUP: H53D23009070001.

## Data availability

The datasets generated for this study are available on request to the corresponding author.

## Notes

### Competing Interest Statement

The authors have declared no competing interest.

